# Mild reductions in guard cell *sucrose synthase* 2 expression leads to slower stomatal opening and decreased whole plant transpiration in tobacco

**DOI:** 10.1101/2020.09.11.293555

**Authors:** Francisco Bruno S. Freire, Ricardo L. G. Bastos, Raissa S. C. Bret, Silvio A. Cândido-Sobrinho, David B. Medeiros, Werner C. Antunes, Alisdair R. Fernie, Danilo M. Daloso

**Author notes:** This work was funded by the National Council for Scientific and Technological Development (CNPq, Grant 428192/2018-1).

## Abstract

The understanding of the dynamics of stomatal movements has increased substantially through genetic manipulation of plant metabolism either at the whole plant level or specifically in guard cells. However, the regulation of stomatal speediness remains not completely elucidated. Here we shown that reduced expression of guard cell s*ucrose synthase 2* (*NtSUS2*) of *Nicotiana tabacum* L. altered the topology and the connectivity of the guard cell metabolic network and the accumulation of metabolites positively correlated with stomatal speediness during dark-to-light transition. This leads to a slower light-induced stomatal opening, lower steady-state stomatal conductance and a strong reduction (up to 44%) in daily whole plant transpiration in the transgenics, when compared to wild type plants. Furthermore, the transgenic lines transpired more or have lower reduction in whole plant transpiration under short water deficit periods, indicating a higher effective use of water under this condition. Our results collectively suggest that the regulation of stomatal movement and speediness involve a complex modulation of the guard cell metabolic network, in which *NtSUS2* has an important role. The results are discussed on the role of guard cell metabolism for the regulation of both stomatal speediness and whole plant transpiration.

## Introduction

It has been estimated that water demand for agricultural will increase ca. 17% by 2025, mostly due the increase in the average global temperature and the fact that drought episodes will become more frequent according to the predicted climate change scenarios (Dai, 2013; Pennisi, 2008; Rahmstorf & Coumou, 2011). It is thus important to improve plant water use efficiency (WUE), defined as the ratio between the amount of accumulated biomass per unit of water used or transpired (Condon et al., 2004). However, plant responses to adverse conditions are modulated by complex regulatory networks, which act at different spatial and temporal scales. This highlights the complexity of plant cell functioning and the difficulty in finding biotechnological targets for plant WUE improvement (Bertolli, Mazzafera & Souza, 2014). One important strategy to improve WUE is decreasing plant water consumption by genetic manipulation of key regulator(s) of stomatal movements (Flexas, 2016; Flütsch et al., 2020b; Gago et al., 2014; McAusland et al., 2016; Papanatsiou et al., 2019). The WUE fundamentally depends on the ratio between photosynthetic carbon assimilation and water lost by the transpiration process, it is reasonable to assume that stomata act as a master regulator of WUE (Brodribb, Sussmilch, & McAdam, 2019). However, although the stomatal development is relatively well understood (Dow & Bergmann, 2014; Qi & Torii, 2018), knowledge concerning the regulation of guard cell metabolism is insufficient, despite this being a great potential target for plant WUE improvement (Daloso et al., 2017; Gago et al., 2020; Lawson & Matthews, 2020).

Stomata are leaf epidermal structures consisting of two guard cells that surround a pore and, in certain cases, with additional subsidiary cells (Lima et al., 2018) whose aperture are actively regulated (Schroeder et al., 2001). Guard cells are highly specialized and integrate endogenous and environmental signals to regulate stomatal opening (Sussmilch, Schultz et al., 2019). Environmental cues such as temperature, soil water status, light, CO_2_ concentration and air vapor pressure deficit modulate stomatal movements in a mesophyll cells-dependent manner (Lawson et al., 2014; Mott, 2009). The dynamics of stomatal movements are thus closely linked to the mesophyll photosynthetic activity, in which the transport of mesophyll-derived metabolites such as sucrose and malate and their import into guard cells seem to be key for stomatal movement regulation (Daloso, dos Anjos, & Fernie, 2016; Gago et al., 2016; Lima et al., 2019; Wang et al., 2019). Indeed, genetic manipulation of genes regulating the trade-off between photosynthetic rate (*A*) and stomatal conductance (*g*_s_) has been shown to be an effective strategy to improve photosynthesis, WUE and/or drought tolerance (Antunes et al., 2017; Araújo et al., 2011; Daloso et al., 2016b; Kelly et al., 2019; Laporte, Shen & Tarczynski, 2002; Lugassi et al., 2015; Nunes-Nesi et al., 2007). Guard cell genetic manipulation has been achieved through the use of guard cell specific promoters such as KST1 (Kelly et al., 2017; Kopka, Provart & Muller-Rober, 1997; Plesch, Ehrhardt & Mueller-Roeber, 2001), which is important to avoid undesired pleotropic modifications in mesophyll cells or sink tissues, notably when sugar-related genes are manipulated.

Several studies indicate the importance of carbohydrate metabolism for the regulation of stomatal movements (Daloso et al., 2016a; Granot & Kelly, 2019; Lima et al., 2018). It has been demonstrated that transgenic plants with modified guard cell sugar metabolism have altered stomatal movements. For instance, transgenic plants with increased expression of *hexokinase* or antisense inhibition of a sucrose transporter (SUT1) have increased WUE (Antunes et al., 2017; Kelly et al., 2019). By contrast, overexpression of *sucrose synthase 3* (*StSUS3*) increased *g*_s_, *A* and plant growth (Daloso et al., 2016b). Additionally, Arabidopsis plants lacking hexose transporters (STP1 and STP4) or enzymes related to starch degradation (AMY3 and BAM1) have altered guard cell sugar metabolism and reduced speed of light-induced stomatal opening (Flütsch et al., 2020a,b). These studies demonstrated that genetic manipulation of guard cell sucrose metabolism is a promising strategy to improve WUE.

Furthermore, the role of sucrose in the regulation of stomatal movements has been reinterpreted on the basis of recent results. These include the demonstration that sucrose can induce stomatal closure in an ABA-mediated, hexokinase-dependent mechanism (Kelly et al., 2013; Lugassi et al., 2015), and that the degradation of sucrose within the guard cells is an important source of substrate for the TCA cycle and glutamine biosynthesis during light-induced stomatal opening (Daloso et al., 2015; Medeiros et al., 2018; Robaina-Estévez et al., 2017). Thus, guard cell sucrose metabolism seems to play a major role in regulating the *A*-*g*_s_ trade-off during both stomatal opening and closure (Granot & Kelly, 2019; Lima et al., 2018).

Sucrose metabolism is not only important for the guard cell but also for the overall carbon distribution throughout the plant. In the cytosol of plant cells, sucrose is degraded into hexoses by different invertase (INV) and sucrose synthase (SUS) isoforms (Fettke and Fernie 2015). The number and the expression of SUS isoforms vary among plant species and organs (Angeles-Núñez & Tiessen, 2012; Baroja-Fernández et al., 2012; Bieniawska et al., 2007; Koch et al., 1992; Kopka et al., 1997). In *Nicotiana tabacum* L., there are seven SUS isoforms (*NtSUS1-7*) and isoforms 2 and 3 are the most abundant in mature leaves (Wang et al., 2015). In *Arabidopsis thaliana* L., recent results indicate that *AtSUS3* is solely expressed in embryo and guard cells (Yao, Gonzales-Vigil & Mansfield, 2020), similar to the expression pattern observed for its ortholog in *Solanum tuberosum* L. (Kopka et al., 1997). Furthermore, guard cell SUS activity is approximately 40-fold higher compared to that of whole leaves (Daloso et al., 2015). Taken together, these data suggest a central role of SUS in the regulation of guard cell metabolism and stomatal movements. Here we show that tobacco transgenic plants with mild reductions in guard cell *NtSUS2* expression exhibited up to 44% less whole plant transpiration than wild type plants, yet only a minor impact on biomass production, corresponding to increased yield WUE (_y_WUE) in one of the transgenic lines under well-watered conditions. Surprisingly, the transgenic lines transpired more under water restriction periods, indicating a more efficient use of water under this condition. Our results are collectively discussed in terms of the role of *NtSUS2* and guard cell sucrose metabolism in the regulation of stomatal movements and whole plant transpiration.

## Materials and methods

### Plant material and growth conditions

*Nicotiana tabacum* L. cv Havana 425 wild type (WT) and transgenic plants in which the *sucrose synthase 2* (*NtSUS2*) gene expression was suppressed under the control of the KST1 promoter (X79779). The transformation was carried out by cloning the *SUS3* gene from *Solanum tuberosum* (*StSUS3*) (STU24088) into the pBinK plasmid vector, which was derived from pBinAR-Kan (Höfgen & Willmitzer, 1990) but had the CaMV-35S promoter replaced by the KST1 promoter. A 1567 bp fragment was obtained by the digestion of *SUS3* gene using the *Kpn*I restriction enzyme (Antunes et al., 2012), which was cloned in the antisense direction in the pBinK vector, between the KST1 promoter and the OCS terminator (Figure 1A). This construct was inserted into *Agrobacterium tumefasciens* (Strain GV 3101) by electroporation and cultivated in suspension with leaf discs from tobacco in order to achieve plant transformation. The transformation was confirmed by PCR of the *NPTII* marker gene that confers kanamycin resistance.

**Figure 1.**
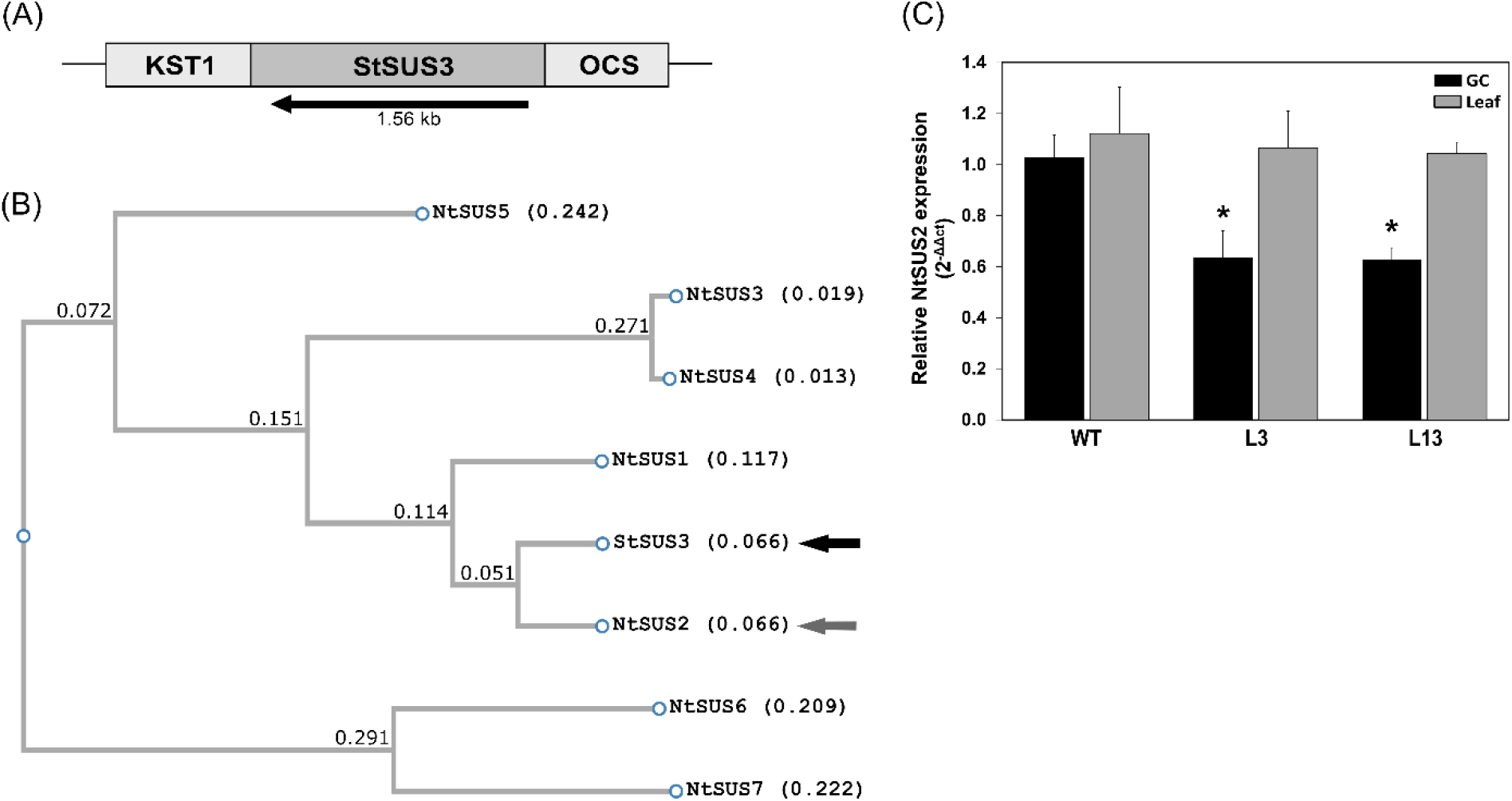
Genetic and phylogenetic characteristics of sucrose synthase (SUS) genes of *Nicotiana tabacum* L. transgenic lines. (A) Simplified scheme of the construction used in this study, in which *Solanum tuberosum* L. *StSUS3* isoform was inserted into tobacco plants in the antisense direction between the KST1 promoter and OCS terminator. (B) Phylogenetic tree of seven tobacco SUS isoforms (*NtSUS1-7*) (*NtSUS2*, grey arrow) and *StSUS3* (black arrow). The numbers refer to the branch length. Similar values between two isoforms indicate that they are genetically closed to each other. (C) Expression of *NtSUS2* gene in leaves and guard cells of wild type (WT) and transgenic lines (L3 and L13) plants. *NtSUS2* gene expression was analyzed by qRT-PCR normalized by protein phosphatase 2A (*PP2A*) gene expression as internal control. Data are shown as relative expression normalized to WT (n = 3 ± SE). Asterisks (*) indicate significant difference from WT by Student’s *t* test at 5% of probability (*P* < 0.05).

Transgenic seeds of the lines L3 and L13 of T_3_ generation were obtained and germinated *in vitro*. These lines were chosen based on preliminary results of transpiration, leaf temperature and gas exchange analysis, as previously reported (Antunes et al., 2017; Daloso et al., 2016b). After sterilization the seeds were germinated in Petri dishes containing Murashige and Skoog (MS) medium (Murashige & Skoog, 1962) with addition of 50 µM of Kanamycin® (KAN) for the transgenic lines and cultivated *in vitro* for 30 days under growth chamber conditions (16h of photoperiod, 100 µmol photons m^-2^ s^-1^, 25 ± 1 °C and relative humidity 53 ± 5%. Seedlings showing KAN resistance were transferred to 0.1 L pots with substrate composed by a mixture of vermiculite, sand and soil (1:1:1) and kept well-watered for 15 days under greenhouse conditions with natural 12 h photoperiod (maximum of 500 µmol photons m^-2^ s^-1^, 30 ± 4 °C and relative humidity 62 ± 10%). Five different experiments were performed in plants growing in soil or hydroponic system under greenhouse condition or in soil under growth chamber condition. The hydroponic experiments were carried out using Hoagland and Arnon nutritive solution in 2.5 L pots (Hoagland & Arnon, 1950). These plants were used for isolation of guard cell-enriched epidermal fragments for metabolomics analysis. Plants growing on soil were cultivated in 2.5 L (for growth chamber) or 5.0 L (for greenhouse) pots containing the same substrate mentioned above. These plants were irrigated with Hoagland and Arnon nutritive solution three times per week. All plants were cultivated during 45 or 60 days until the beginning of each experiment. A water deficit experiment was performed by suspending irrigation on 45 day-old plants cultivated under growth chamber conditions.

### Phylogenetic and gene expression analyses

Multiple sequence alignment was performed using ClustalW software (Kyoto University Bioinformatics Center, Kyoto, Japan) utilizing CDS of *NtSUS1-7* and *StSUS3* to build a phylogenetic tree by FastTree tool (Price, Dehal, and Arkin 2009). Total RNA from leaves and guard cells of 45 days-old plants were isolated and the cDNA synthesized using SV Total RNA Isolation System and M-MLV Reverse Transcriptase (Promega Corporation, Madison, Wisconsin, EUA). The primer for protein phosphatase 2A (*PP2A*) was designed by aligning the coding sequence (CDS) obtained from NCBI database using the muscle tool in Mega X software (FW 5’-CACTTCAGTCAATTGATAACGTC-3’; REV 5’-GCAAAATCCTACCAAAGAGGG-3’). The primers used to investigate *NtSUS* expression were obtained from previous work (Wang et al., 2015) (FW 5’-CACATTGATCCATACCACGGGGAT-3’; REV 5’-ACAGCAGCCAGTGTCAACAACCGA-3’). In order to estimate *SUS* silencing, qRT-PCR was performed by using GoTaq® qPCR Master Mix (Promega Corporation, Madison, Wisconsin, EUA) accordingly to the manufacturer instructions. Relative transcripts expression was calculated by the 2^-ΔΔCt^ method, in which *PP2A* was used as internal control and WT was used as calibrator (Livak & Schmittgen, 2001).

### Photosynthetic light and CO_2_ response curves

Photosynthetic response curves to light (*A*-PAR), CO_2_ (*A*-*C*_i_) were performed in completely expanded leaves from 45 day-old plants of all genotypes using a portable infrared gas exchange analyser (IRGA) (LiCor 6400XT, Lincoln, NE, USA). *A*-PAR curves were measured using 400 ppm CO_2_, block temperature at 28 °C and 10% of blue light, whilst *A*-*C*_i_ were carried out using 1000 µmol photons m^-2^ s^-1^. A three component exponential rise to maximum equation was used to fit the photosynthetic curves: *A* = *a* (1 - *e*^*-bx*^) + *c*, where *A* = photosynthetic rate, x = PAR or substomatal CO_2_ concentration (*C*_i_), and a, b, c are parameters estimated by the non-linear regression (Watling, Press & Quick, 2000).

### Whole plant transpiration (WPT) and growth analysis

WPT was determined by a gravimetric methodology (Daloso et al., 2016b). Soil-filled pots without plants were used to estimate direct evaporation from soil. The pots were irrigated with water at beginning of the night in a daily basis, except at the days in which the water was withdrawn, as indicated in the figures. The pots were weighed at predawn and at the end of the days. The daily WPT (g H_2_O d^-1^ plant^-1^) was obtained by the difference between the two weights and by subtracting the evaporation. WPT was further recorded between different time intervals of a daily course. Leaves larger than 5 cm length were used to estimate leaf area (LA) by a previously described model (Antunes et al., 2008, 2017). LA was recorded in three different days of the experiments, whilst the days in between LA was estimated by linear regression (r^2^ > 0.98). The LA (cm^2^) was used to calculate specific leaf area (cm^2^ g^-1^ leaf dry weight (DW)) as well as to estimate WPT per leaf area (g H_2_O d^-1^ m^-2^). At the end of WPT experiments, it was determined the total leaf number (> 5cm length), stem length and the DW of leaf, stem and roots by drying it at 80 °C for seven days. These parameters were used to determined total dry biomass (leaf + stem + roots), harvest index (leaf /total biomass), shoot DW (leaf + stem), shoot/root (g g^-1^), leaves/root (g g^-1^) and plant leaf area/roots (cm^2^ g^-1^). The relative growth rate (RGR, g g^-1^ day^-1^) was obtained by the equation: RGR = ln (final weight) – ln (initial weight)/final day – initial day following previous study (Hoffmann and Poorter 2002), in which the initial DW of leaf and total biomass were estimated by regression using leaf area measurements (Figure S1).

### Determination of plant water use efficiency (WUE)

Intrinsic WUE (_i_WUE – *A*/*g*_s_ ratio) (µmol CO_2_ mmol^-1^ H_2_O) was estimated from steady state values of *A* and *g*_s_ under 1000 µmol photons m^-2^ s^-1^ and 400 ppm CO_2_ from *A*-PAR curves and stomatal opening kinetics (described below). The data from WPT and biomass were used to estimate season-long WUE (_sl_WUE - ratio between the amount of accumulated biomass per unit of water transpired) and yield WUE (_y_WUE - ratio between the yield of the harvestable organ (leaves) to water transpired over time) (Bacon 2004). For the determination of both _y_WUE (g DW leaves kg^-1^ H_2_O transpired) and _sl_WUE (g DW kg^-1^ H_2_O transpired), we used LA data to estimate the leaf and total biomass DW of the beginning of the experiment by linear regression with accuracy of R^2^ > 0.82 for leaves and R^2^ > 0.95 for total biomass (Figure S1).

### Stomatal opening kinetics during dark-to-light transition

Stomatal opening kinetics were measured in fully expanded, dark-adapted leaves. In order to avoid circadian effects in the measurements, only three plants were measured at a time. All gas exchange parameters were recorded every 10 sec for 300 sec in the dark plus 3200 sec under 1000 µmol photons m^-2^ s^-1^, 400 ppm CO_2_ and block temperature 28 °C using an IRGA. The curves were plotted by averaging 10 readings every 100 sec. The half-time needed to *g*_s_ reach steady state (t_50%_) were estimated using the *g*_s_ curves (Lima et al., 2019). The maximum slope of *g*_s_ response (*Sl*_max_) was estimated by calculating d*g*_s_/dt when the rates of change reached an acceleration plateau. Non-linear regression equations were obtained (*y* ∼ *y*_o_ + *a* (1 - *e* ^(-*bx*)^)) and further derived to obtain the rate of change. Subsequently, the data undergone curve fitting using linear plateau model (*y* ∼ *a* + *b* (*x* - *c*) (*x* ≤ *c*)) to estimate the time when *g*_s_ reach maximum rate of change in light-induced stomatal opening (Figure S2).

### Stomatal density and fresh weight loss in detached leaves

Determination of stomatal density (SD) (number of stomata mm^-2^) of abaxial and adaxial leaf surfaces and fresh weight loss in detached leaves were determined in 45 day-old plants as previously described (Daloso et al., 2016b).

### Isolation of guard cell-enriched epidermal fragments and metabolite profiling analysis

Guard cell-enriched epidermal fragments (after simply called guard cells) were isolated as described earlier (Daloso et al., 2015). Guard cells were harvested at pre-dawn and in the early morning (120 min after sunrise) and used for metabolomics analysis. Metabolite extractions were performed using ∼200 mg of guard cells. Polar metabolites were extracted using a well-established gas chromatography coupled to *time of flight* mass spectrometry (GC-TOF-MS) platform (Lisec et al., 2006). Chromatogram and mass spectral analyses were carried out using TagFinder software (Luedemann et al., 2008).

### Metabolic network analysis

Correlation-based networks were created between relative guard cell metabolite content in dark and light for each genotype and between stomatal speediness parameters (*Sl*_max_ and t_50%_) and relative guard cell metabolite changes in the dark and after the transition to the light using all genotypes data, in which the nodes correspond to the metabolites and the links to the strength of connection between nodes in module (positive or negative) by Pearson correlation. The relative guard cell metabolic changes were obtained by dividing the content of each metabolite in the light *per* the average of those found in the dark within the genotype. The networks were designed by restricting the strength of the connections to a specific limit of Pearson correlation coefficient (*r*) (−0.5 > *r* > 0.5). The network parameters, clustering coefficient, network heterogeneity, network density, network centralization and connected components were obtained as described in previous work (Assenov et al., 2008). Preferential attachment is a characteristic of scale-free networks in which as higher is the number of links of a node, higher is the probability of this node to receive new links (Albert & Barabási, 2002). Here, we determined the preferential attachment as the nodes that are considered as hub in the dark and that maintained the number of links in the light higher than the average of the network in the dark.

### Statistical analysis

The transgenic lines were statistically compared to WT using a Student’s *t* test at 5% of probability (*P* < 0.05) by using Microsoft Excel (Microsoft, Redmond, WA, USA). Regression analysis were carried out using SIGMAPLOT 14 (Systat Software Inc., San Jose, CA, USA). The equations from stomatal kinetics regressions were derived and the rate of change were fit to linear plateau using easyreg package in R 3.6.3 (Arnhold, 2018; R Core Team, 2020). Correlation analysis were carried out by Pearson correlation analysis using the Java-based CorrelationCalculator software (Basu et al., 2017). The metabolomics data were analyzed using the MetaboAnalyst platform (Chong et al., 2018). Multivariate analysis such as partial least square-discriminant analysis (PLS-DA) and orthogonal PLS-DA (orthoPLS-DA) was performed in Cube root-transformed and mean-centered data by using both Cube root and Pareto-scaling mode of the MetaboAnalyst platform, which is recommended to reduce the scale variability of metabolomics datasets (Xia & Wishart, 2011). Correlation-based metabolic networks were designed by using MetScape on CYTOSCAPE v.3.7.2 software (Karnovsky et al., 2012; Shannon et al., 2003).

## Results

### Phylogenetic analysis indicates that NtSUS2 is ortholog of StSUS3

We have previously cloned potato *sucrose synthase 3* (*StSUS3*) and inserted in the sense direction into tobacco guard cells in order to investigate the function of this gene in guard cell metabolism and stomatal movements (Daloso et al., 2016b). The reasons to use *StSUS3* and tobacco plants are based in the fact that *StSUS3* and its ortholog *AtSUS3* has been previously shown to be highly expressed in guard cells (Bates et al., 2012; Bauer et al., 2013; Kopka et al., 1997; Yao et al., 2020) and that tobacco produce large leaves, which is crucial to harvest sufficient guard cells for metabolomics analysis (Daloso et al., 2015). Here, *StSUS3* was inserted in the antisense orientation into tobacco leaves under control of the KST1 promoter (Figure 1A). In order to identify which tobacco SUS (*NtSUS*) isoform corresponds to *StSUS3*, we first aligned all seven *SUS* CDS sequences from tobacco with *StSUS3*. The phylogenetic tree confirmed previous study (Wang et al., 2015) indicating that *StSUS3* is ortholog of *NtSUS2*. (Figure 1B). Here, after we refer to *NtSUS2* expression. In fact, qRT-PCR analysis showed that the *NtSUS2* expression decreased by almost 40% in guard cells of L3 and L13 lines compared to wild type (WT), whilst no difference in leaf gene expression was observed (Figure 1C). Although *StSUS3* is also similar to *NtSUS1* (Figure 1B), we did not detect the expression of this isoform in guard cells and leaves, in agreement with a previous study (Wang et al., 2015).

### Gas exchange analysis in plants with reduced guard cell NtSUS2 expression

Only L3 displayed lower *A* compared to WT in either light or CO_2_ response curves determined in well-watered plants (Figures 2A,B). The data from these curves also revealed that no differences in the steady state values of transpiration rate (*E*), dark respiration (R_d_) and intrinsic WUE (iWUE) (*A*/*g*_s_ ratio) were apparent between WT and transgenic lines, whilst L3 and L13 displayed reduced steady state stomatal conductance (*g*_s_) compared to WT plants (Figures 2C-F). We also evaluated the kinetics of gas exchange during the dark-to-light transition (Figures 3, S2). No statistical difference was observed in *A* between WT and transgenic lines (Figure 3A). The light stomatal response was, in general, slower in the transgenic lines, which alter the dynamic of both _i_WUE and *C*_i_/*C*_a_ and leads to significant reductions in *Sl*_max_ in L3 and t_50%_ in both L3 and L13 (Figures 3B-F). Interestingly, the dynamic of WT *g*_s_ curves have a decrease in *g*_s_ after 2000 sec, and this was not observed in the transgenic lines (Figure 3B).

**Figure 2.**
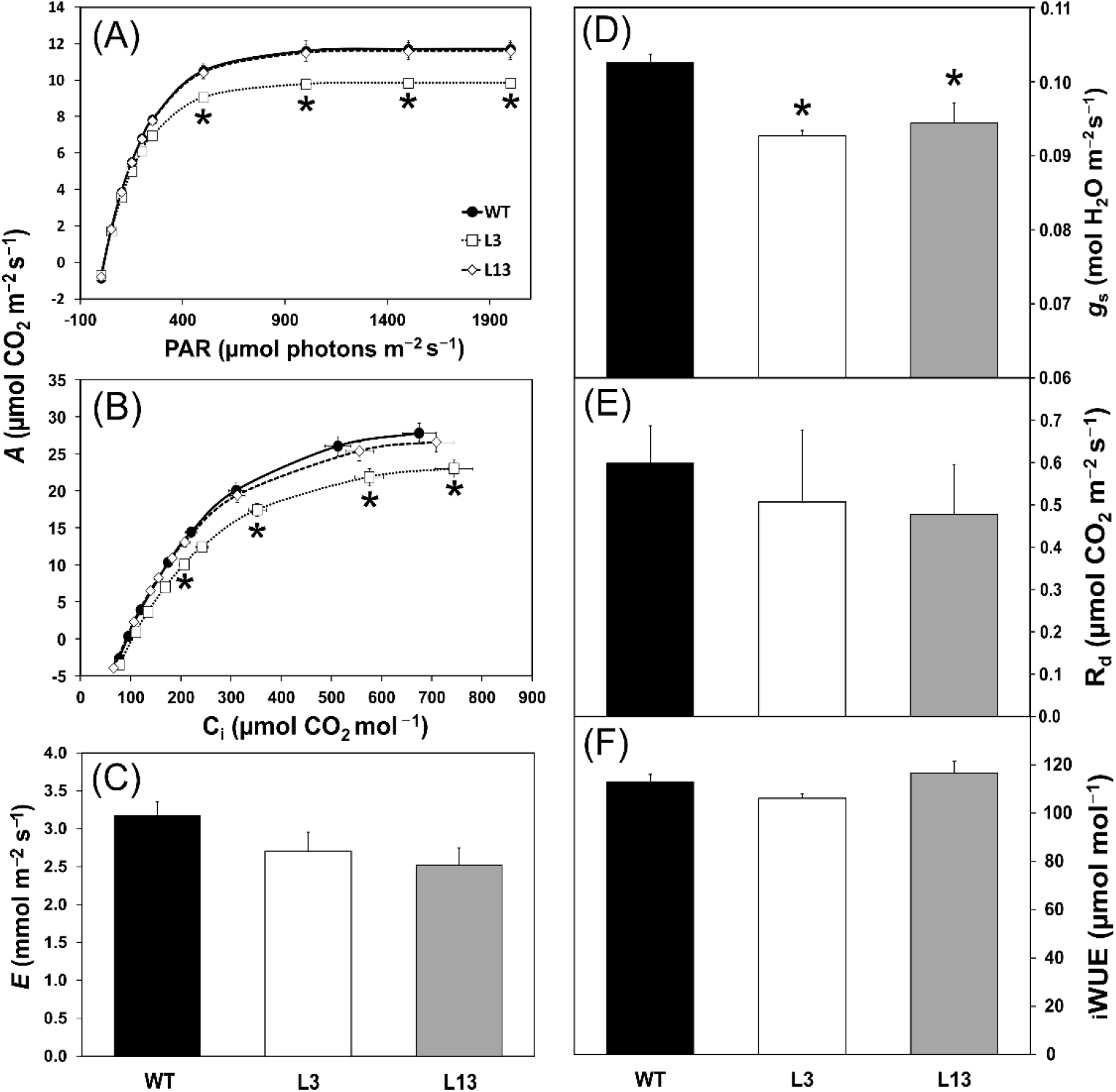
Physiological characterization of *Nicotiana tabacum* L. wild type (WT) and transgenic lines (L3 and L13) antisense to *NtSUS2* grown under well-watered conditions. (A-B) Photosynthetic response curves to light (*A*-PAR) and substomatal CO_2_ concentration (*A*-*C*_i_). The regression line was determined using the equation *A* = *a* (1 − *e*^*-bx*^) + *c*. (C-F) Steady state values of transpiration rate (*E*), stomatal conductance (*g*_s_), dark respiration (R_d_) and water use efficiency (_i_WUE) (*A*/*g*_s_) from gas exchange analysis. Measurements were taken in plants grown under greenhouse and well-watered conditions. Asterisks (*) indicate significant difference from WT by Student’s *t* test at 5% of probability (*P* < 0.05). (n = 4 ± SE).

**Figure 3.**
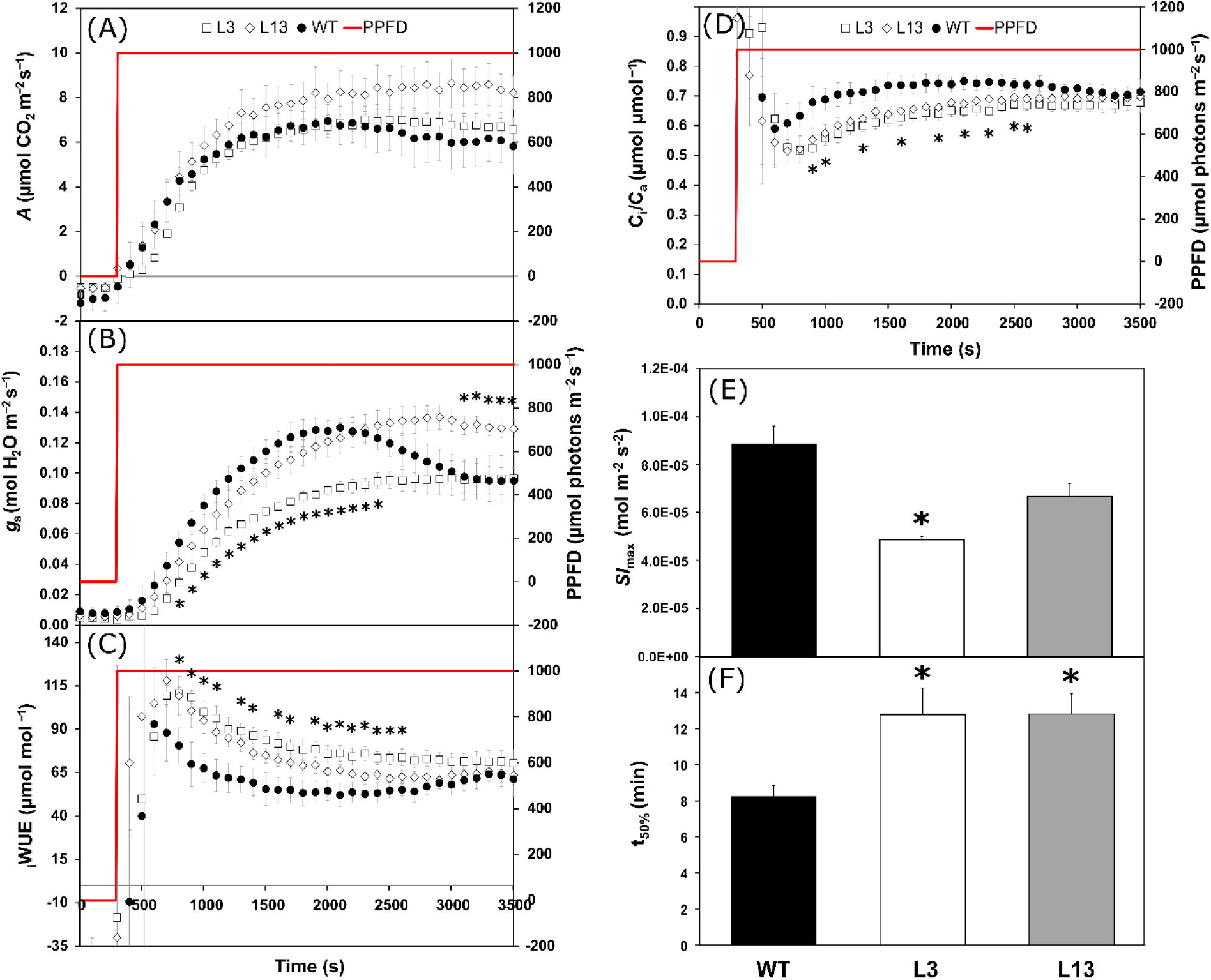
Kinetic curves of physiological parameters and speediness of stomatal opening during dark-to-light transition of *Nicotiana tabacum* L. wild type (WT) and transgenic lines (L3 and L13) antisense to *NtSUS2* grown under growth chamber and after water deficit stress. (A-D) Kinetic of photosynthetic rate (*A*), stomatal conductance (*g*_s_), intrinsic water use efficiency (_i_WUE) and the ratio of substomatal and ambient CO_2_ concentration (*C*_i_/*C*_a_). (E) Maximum slope (*Sl*_max_) parameters obtained through the linear plateau model using the time when *g*_s_ reach maximum rate of change in light-induced stomatal opening. (F) Half-time (t_50%_) needed to *g*_s_ reach steady state in light-induced stomatal opening. Measurements were taken in plants grown under growth chamber and after water deficit (n = 3 ± SE). Asterisks (*) indicate significant difference from WT by Student’s *t* test at 5% of probability (*P* < 0.05).

### Mild reductions in guard cell NtSUS2 expression substantially reduce whole plant transpiration (WPT) under well-watered and fluctuating environmental conditions

We next used a gravimetric methodology to determine WPT in plants grown under well-watered (WW) and non-controlled greenhouse conditions, which varies substantially in terms of light, temperature and humidity throughout the days (Figure 4A). Under these conditions, both L3 and L13 showed significantly lower WPT (up to 44%) compared to WT throughout the eleven days of the experiment (Figures 4B,C). At day 5, plants reached a peak of WPT (Figure 4B), which was associated to a sequence of sunny days (Figure 4A). Following the interruption of irrigation at day 5, a strong reduction in WPT in WT plants was observed at day 6, whilst no changes in WPT of the transgenic lines were observed (Figure 4B). On this day, the percentage of WPT was invariant across the genotypes (Figure 4C). When the cumulative WPT across the experiment were compared, L3 and L13 were found to transpire approximately 32% less than the WT (Figure 4D), meaning that they respectively conserved 416 and 433 g H_2_O plant^-1^ during the experiment.

**Figure 4.**
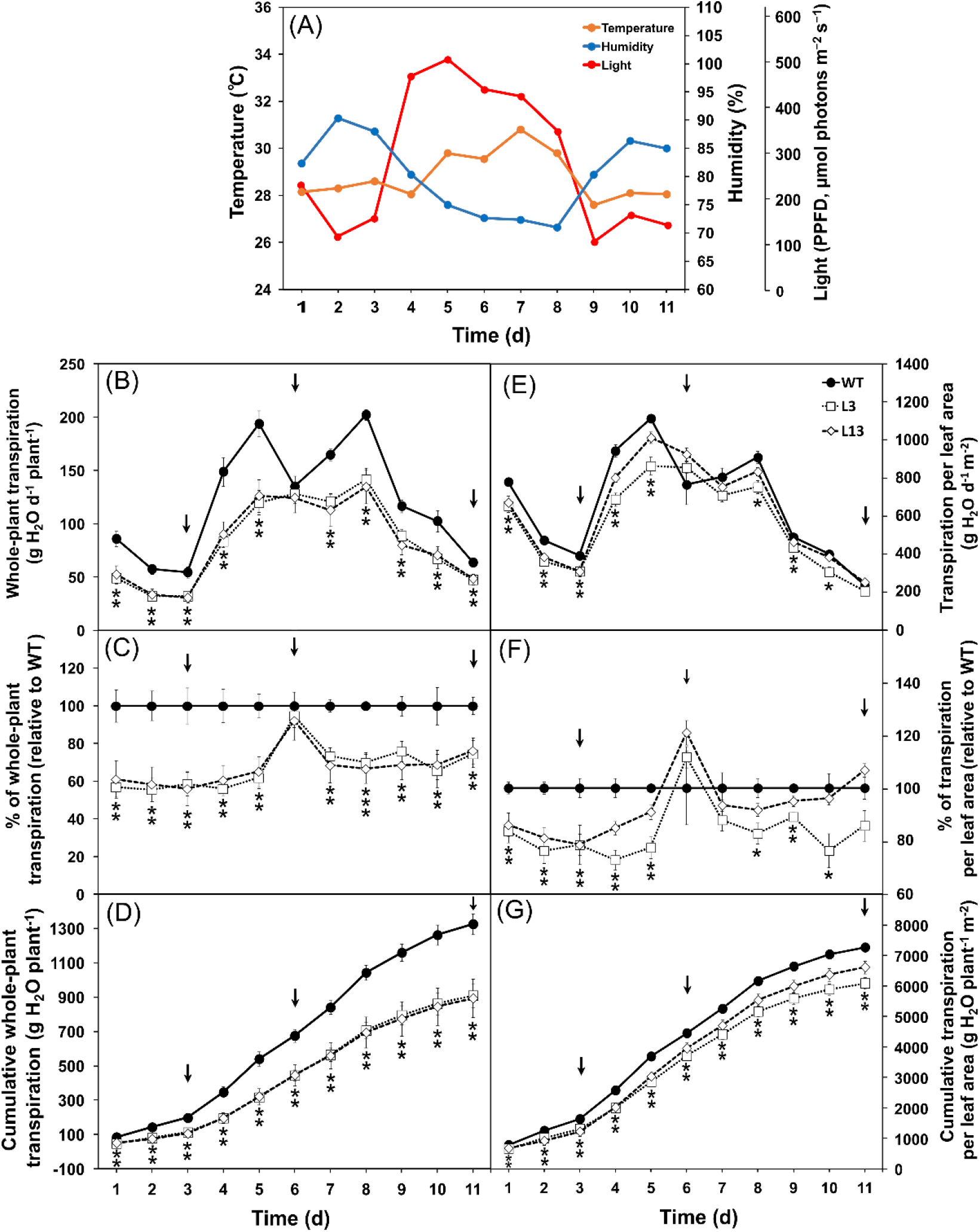
Whole-plant transpiration (WPT) of *Nicotiana tabacum* L. wild type (WT) and transgenic lines (L3 and L13) antisense to *NtSUS2* grown under greenhouse and well-watered conditions. (A) Daily average of light, temperature and humidity throughout eleven days of the whole-plant transpiration experiment carried out under greenhouse conditions. (B and E) Daily WPT per plant (g H_2_O day^-1^ plant^-1^) (B) or per leaf area (mg H_2_O day^-1^ cm^-2^) (E). (C and F). % of daily WPT per plant (C) or per leaf area (F) relative to WT. (D and G) Accumulated water loss per plant (D) or leaf area (G) throughout the experiment. Black arrows indicate days of no watering. One (*) and two asterisks (**) indicate that one or two transgenic lines are significant different from WT by Student’s *t* test at 5% of probability (*P* < 0.05), respectively. (n = 5 ± SE).

The leaf area was initially smaller in L3 and L13 plants compared to WT. However, from day 5 to 11 of the experiment, only the leaf area of line L13 remained smaller than WT (Figure S3). WPT per leaf area unit (g H_2_O m^-2^ d^-1^) was also lower in L3 (28%) and L13 (22%) than WT (Figures 4E,F), which lead to a lower cumulative water loss per leaf area in these lines (Figure 4G). No changes in leaf stomatal density (SD) and water loss from detached leaves among the genotypes was observed (Figures S4A,B). At the end of the experiment, plants were harvested and growth parameters determined. Although relative growth rate (RGR), total leaf number, specific leaf area and stem length were not significantly altered in the transgenic lines, both L3 and L13 exhibited a reduced total biomass (Table 1). This was associated to a reduced carbon allocation toward the roots, given that reduced roots DW and % of DW roots were observed in these lines (Table 1). By contrast, line L3 showed higher % DW leaf, % DW shoot and shoot-to-root and leaf-to-root ratios than WT (Table 1). It is noteworthy that the leaves are the harvestable part of tobacco plants and the harvest index of L3 and L13 reached 0.58 and 0.54 g g^-1^, but only L3 was significantly higher than WT (0.52 g g^-1^). Line L3 also have higher _y_WUE than WT, whilst no difference in season-long WUE (_sl_WUE) under WW condition was observed (Table 1). Taken all WUE parameters together, reduced expression of the guard cell *NtSUS2* did not alter both steady-state _i_WUE and _sl_WUE, while L3 showed higher _y_WUE and increased _i_WUE in both transgenic lines during dark-to-light transition was observed.

**Table 1.**
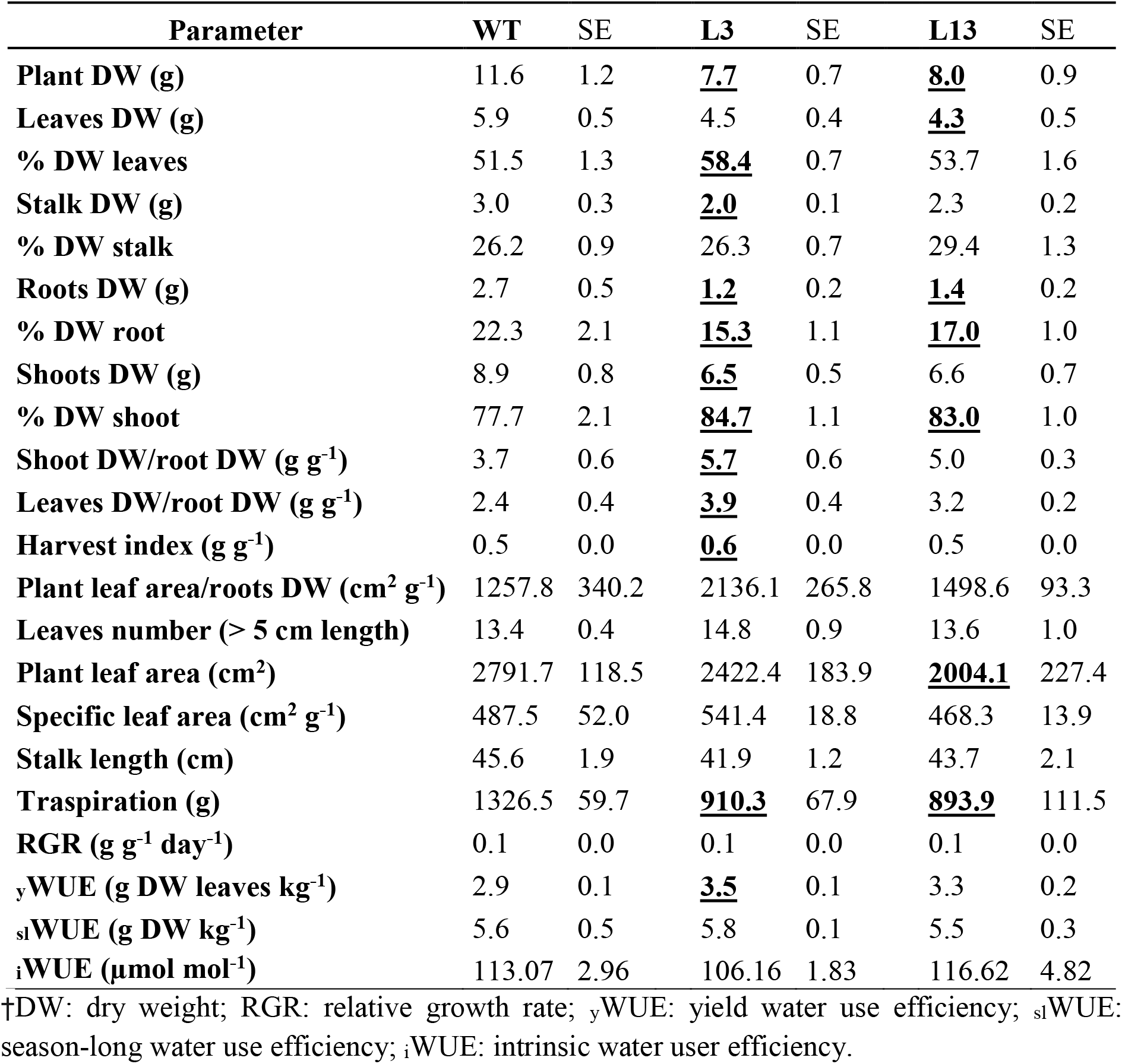
Growth parameters of 60 days-old *Nicotiana tabacum* L. wild type (WT) and *NtSUS2* antisense transgenic (L3 and L13) plants grown under greenhouse and well-watered conditions. Average values in bold and underlined type are significant different from WT by Student’s *t* test at 5% of probability (*P* < 0.05). (n = 5 ± SE).

| Parameter | WT | SE | L3 | SE | L13 | SE |
| --- | --- | --- | --- | --- | --- | --- |
| Plant DW (g) | 11.6 | 1.2 | <u>**7.7**</u> | 0.7 | <u>**8.0**</u> | 0.9 |
| Leaves DW (g) | 5.9 | 0.5 | 4.5 | 0.4 | <u>**4.3**</u> | 0.5 |
| % DW leaves | 51.5 | 1.3 | <u>**58.4**</u> | 0.7 | 53.7 | 1.6 |
| Stalk DW (g) | 3.0 | 0.3 | <u>**2.0**</u> | 0.1 | 2.3 | 0.2 |
| % DW stalk | 26.2 | 0.9 | 26.3 | 0.7 | 29.4 | 1.3 |
| Roots DW (g) | 2.7 | 0.5 | <u>**1.2**</u> | 0.2 | <u>**1.4**</u> | 0.2 |
| % DW root | 22.3 | 2.1 | <u>**15.3**</u> | 1.1 | <u>**17.0**</u> | 1.0 |
| Shoots DW (g) | 8.9 | 0.8 | <u>**6.5**</u> | 0.5 | 6.6 | 0.7 |
| % DW shoot | 77.7 | 2.1 | <u>**84.7**</u> | 1.1 | <u>**83.0**</u> | 1.0 |
| Shoot DW/root DW (g g<sup>-1</sup>) | 3.7 | 0.6 | <u>**5.7**</u> | 0.6 | 5.0 | 0.3 |
| Leaves DW/root DW (g g<sup>-1</sup>) | 2.4 | 0.4 | <u>**3.9**</u> | 0.4 | 3.2 | 0.2 |
| Harvest index (g g<sup>-1</sup>) | 0.5 | 0.0 | <u>**0.6**</u> | 0.0 | 0.5 | 0.0 |
| Plant leaf area/roots DW (cm<sup>2</sup> g<sup>-1</sup>) | 1257.8 | 340.2 | 2136.1 | 265.8 | 1498.6 | 93.3 |
| Leaves number (> 5 cm length) | 13.4 | 0.4 | 14.8 | 0.9 | 13.6 | 1.0 |
| Plant leaf area (cm<sup>2</sup>) | 2791.7 | 118.5 | 2422.4 | 183.9 | <u>**2004.1**</u> | 227.4 |
| Specific leaf area (cm<sup>2</sup> g<sup>-1</sup>) | 487.5 | 52.0 | 541.4 | 18.8 | 468.3 | 13.9 |
| Stalk length (cm) | 45.6 | 1.9 | 41.9 | 1.2 | 43.7 | 2.1 |
| Transpiration (g) | 1326.5 | 59.7 | <u>**910.3**</u> | 67.9 | <u>**893.9**</u> | 111.5 |
| RGR (g g<sup>-1</sup> day<sup>-1</sup>) | 0.1 | 0.0 | 0.1 | 0.0 | 0.1 | 0.0 |
| <sub>y</sub>WUE (g DW leaves kg<sup>-1</sup>) | 2.9 | 0.1 | <u>**3.5**</u> | 0.1 | 3.3 | 0.2 |
| <sub>sl</sub>WUE (g DW kg<sup>-1</sup>) | 5.6 | 0.5 | 5.8 | 0.1 | 5.5 | 0.3 |
| <sub>i</sub>WUE (μmol mol<sup>-1</sup>) | 113.07 | 2.96 | 106.16 | 1.83 | 116.62 | 4.82 |

†DW: dry weight; RGR: relative growth rate; <sub>y</sub>WUE: yield water use efficiency; <sub>sl</sub>WUE: season-long water use efficiency; <sub>i</sub>WUE: intrinsic water user efficiency.

### WPT under water deficit and growth chamber conditions

Given that both L3 and L13 have demonstrated great differences on water consumption in the days after water withdrawal compared to WT plants, we next subjected a new set of plants to water deficit (WD) under growth chamber conditions (24.5 °C, humidity 55.5% and 100 µmol photons m^-2^ s^-1^ on average). As previously, before the suspension of irrigation, the transgenic lines transpired slightly less than WT plants, with significant difference observed in L13 at the second day of the experiment (Figures 5A,B). However, under WD condition, WT transpiration dropped whilst transgenic lines transpired more (up to 37% in L13) than WT at the day 5, which corresponds to two days after water withdrawn (Figures 5A,B). From day 5 to 8, WPT decreased in all genotypes, but L13 kept transpiring more than WT until day 8 (Figure 5B). Considering the total WPT throughout this experiment, L3 and L13 transpired 19 g H_2_O plant^-1^ (5.4%) and 57 g H_2_O plant^-1^ (16.3%) more than WT, respectively (Figure 5C). Comparing the accumulated WPT of WT and the transgenic lines under WW and WD periods separately, L3 and L13 transpired 7.8 and 15.5% less than WT under WW, respectively, but with no statistical significance. By contrast, L3 and L13 transpired 11.1 and 36.74% more than WT under WD condition, respectively, but only L13 was statistically different from WT (Figure 5C inset).

**Figure 5.**
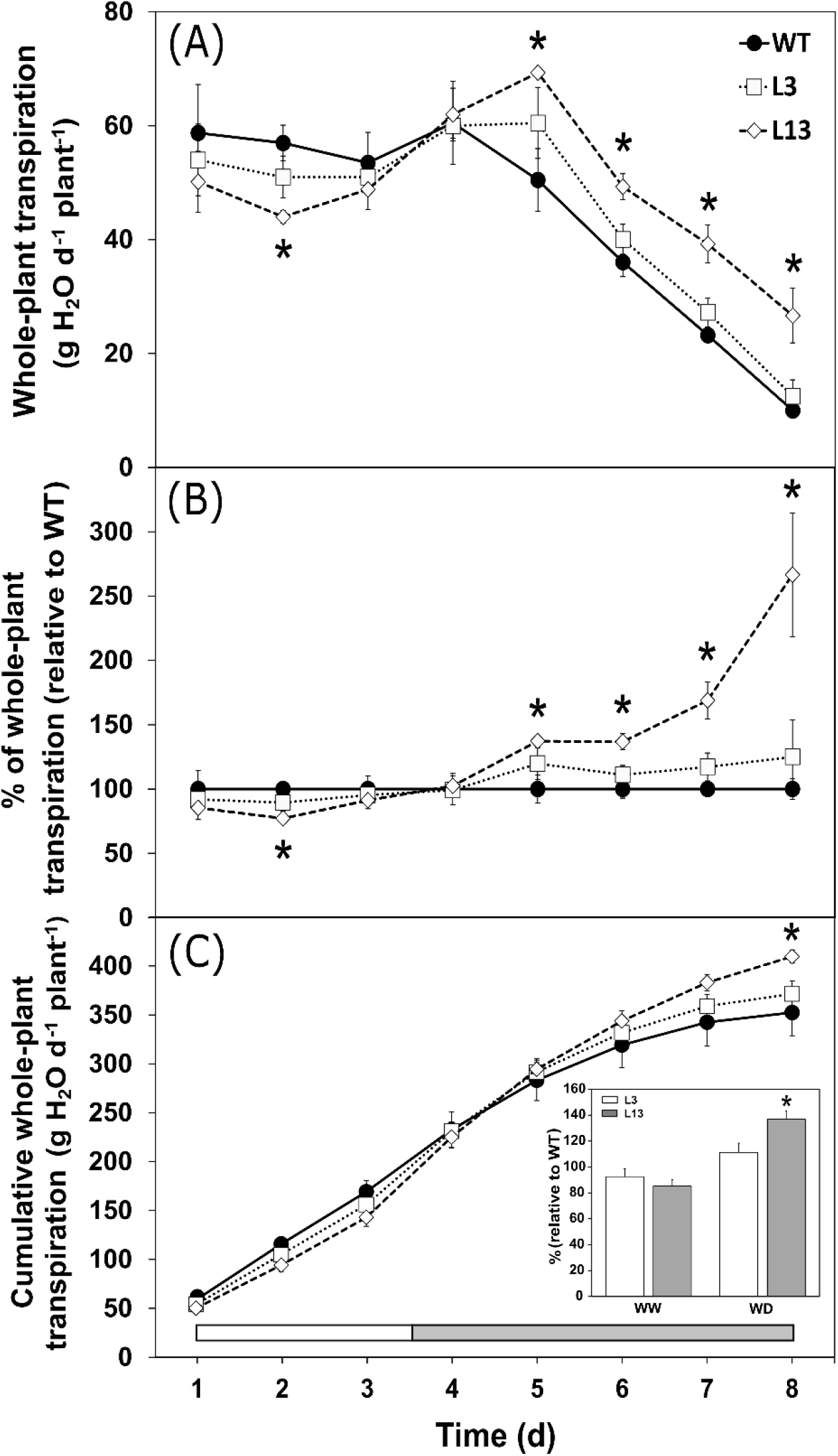
Whole-plant transpiration (WPT) of *Nicotiana tabacum* L. wild type (WT) and transgenic lines (L3 and L13) antisense to *NtSUS2* grown under water deficit stress. (A) Daily WPT per plant (g H_2_O day^-1^ plant^-1^). (B) % of daily WPT per plant relative to WT. (C) Accumulated water loss per plant throughout the experiment. Inset, % of accumulated water loss relative to WT on the first three days of well-watered (WW) and on the next five days of non-watered condition (WD). White and grey bars on the x-axis indicate WW and WD days, respectively. Asterisks (*) indicate significant difference from WT by Student’s *t* test at 5% of probability (*P* < 0.05). (n = 4 ± SE).

### Guard cell metabolic alterations induced by NtSUS2 silencing

We have carried out a metabolite profiling analysis in guard cells of WT, L3 and L13 harvested in the dark and 120 min after the transition to the light. Partial least square discriminant analysis (PLS-DA) and orthogonal PLS-DA (orthoPLS-DA) were performed using the relative metabolic changes observed in each genotype during dark-to-light transition. Both PLS-DA and orthoPLS-DA indicate that reduced guard cell *NtSUS2* expression substantially alter the relative metabolic changes during dark-to-light transition, given that both L3 and L13 were clearly separated from WT by the first components (Figures 6A,B; S5A,B). Scatter plots (S-plots) from the orthoPLS-DA and variable importance in projection (VIP) scores of the PLS-DA models demonstrate that several metabolites pertaining to the groups of carbohydrates, amino acids and tricarboxylic acid (TCA) cycle intermediates were responsible for the discrimination observed among the genotypes (Figures 6C,D; S5C,D). Amino acids such as serine and alanine and the TCA cycle intermediates citrate, isocitrate, aconitate and fumarate have increased relative content whilst adipic acid and trehalose have lower relative content in WT, when compared to the transgenic lines after dark-to-light transition (Figures 6C,D; S5C,D).

**Figure 6.**
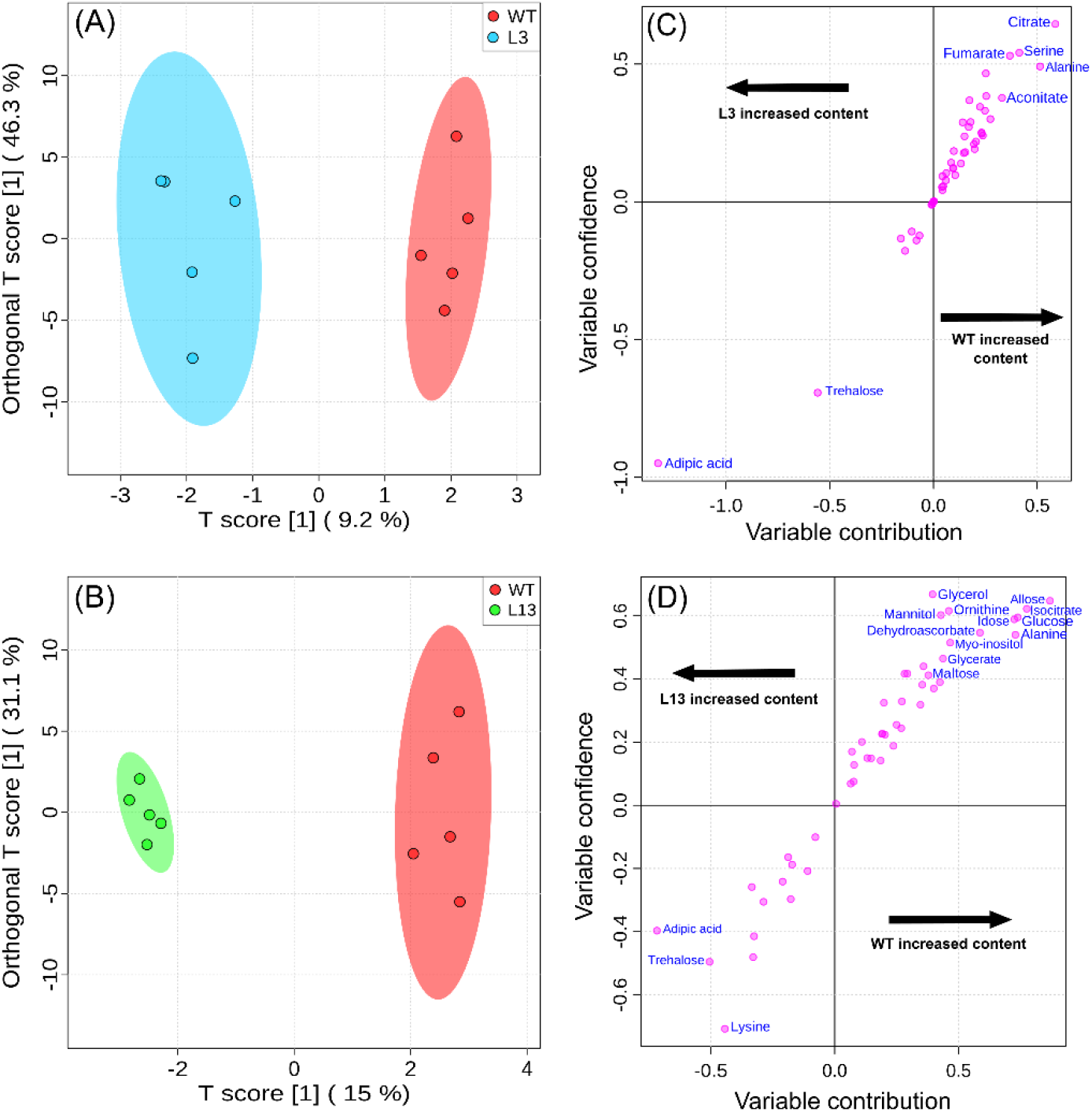
Orthogonal partial least squares-discriminant analysis (orthoPLS-DA) of guard cell metabolomics data. (A-B) OrthoPLS-DA and (C-D) scatter plots (S-plots) of metabolites identified in WT *vs* L3 and L13, separately. Highlighted metabolites in S-plots are the significant ones which have more contribution to the clustering with high variable confidence (y, correlation) in module (−0.4 > y > 0.3 for WT vs L3 and -0.4 > y > 0.4 for WT *vs* L13) and variable importance in projection (VIP) scores of PLS-DA model above 1 (listed in figures S5 C and D). These analyses were carried out using the relative guard cell metabolic changes observed in each genotype in the dark and after the transition to the light. These analyses were performed using MetaboAnalyst platform. (n = 5).

### NtSUS2 silencing alter the topology and the connectivity of the guard cell metabolic network during dark-to-light transition

We next evaluated the effect of *NtSUS2* silencing on guard cell metabolism at metabolic network level, in which metabolites are the nodes and the link is the strength of the connection among them, determined by Pearson correlation coefficient (−0.5 > *r* > 0.5). Correlation-based networks demonstrate that light imposition changed substantially the topology and the connectivity of the networks, with opposite trends observed between WT and the transgenic lines (Figures 7A-F). Guard cell WT metabolic network changed from a highly integrated and connected network under dark condition to a less connected and fragmented network in the light (Figure 7A-B). This is evidenced by the 2.2-fold higher network density, the higher average number of links and the lower number of both connected components and isolated nodes in WT in the dark, when compared to WT in the light (Table S1). Furthermore, WT metabolic network was 1.4-fold and 1.5-fold more connected than L3 and L13 metabolic networks in the dark, respectively. By contrast, L3 and L13 metabolic networks were respectively 1.3-fold and 1.9-fold more connected than WT in the light (Figures 7A-F; Table S1).

**Figure 7.**
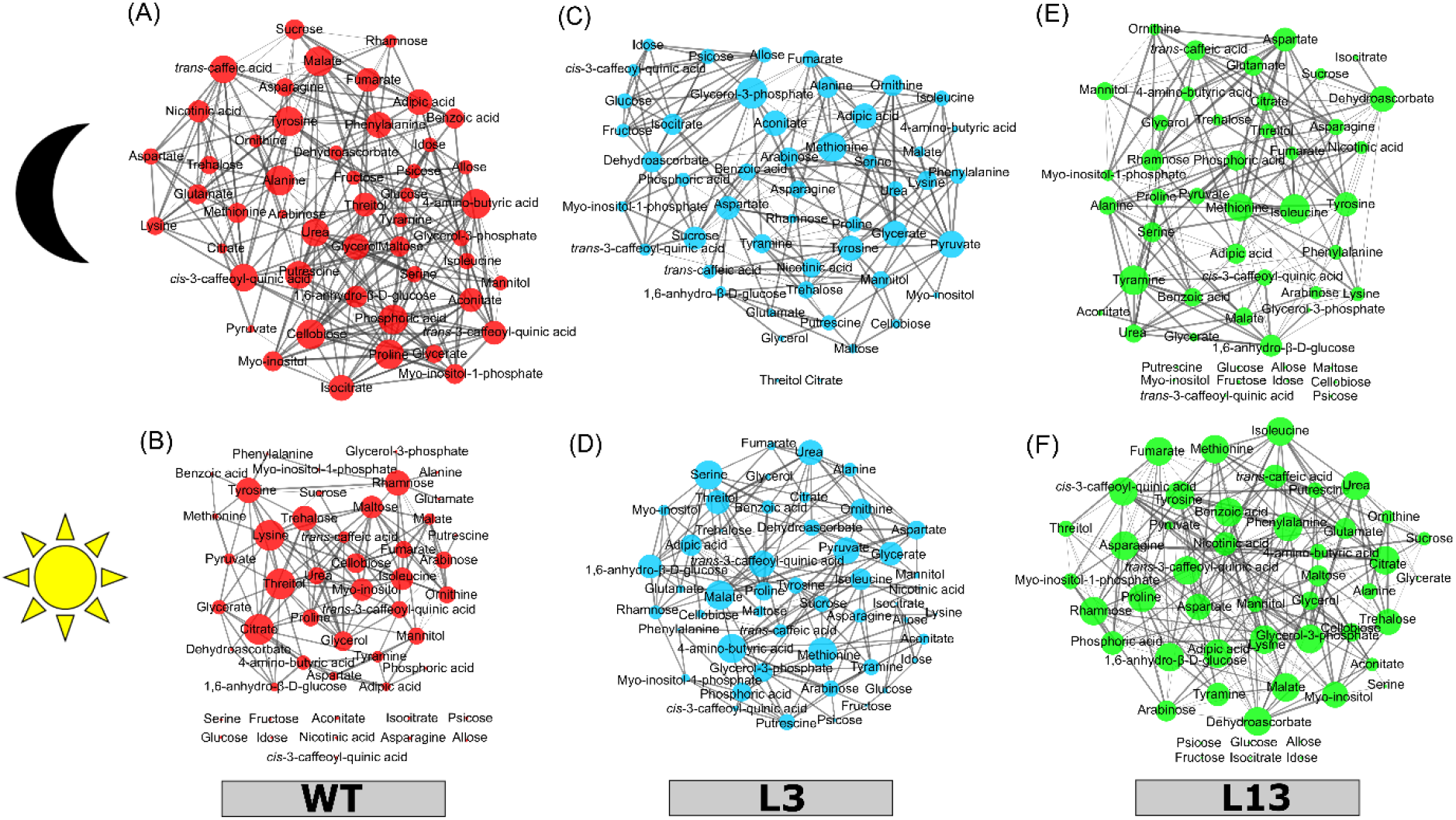
Correlation-based metabolic networks of guard cell metabolomics data from *Nicotiana tabacum* L. wild type (WT) and transgenic lines (L3 and L13) antisense to *NtSUS2* grown under greenhouse and well-watered conditions. The networks were created using data from guard cell metabolite contents observed in each genotype in the dark (A, C, E) and after the transition to the light (B, D, F). Thicker links indicate higher *r*, in module. Bigger nodes indicate higher degree of connection. (n = 5).

Another clear difference observed among WT and transgenic lines is related to the heterogeneity of the network, a topological characteristic of complex networks (Jeong et al., 2000; Pinheiro & Hartmann, 2017). Higher heterogeneity value reflects the tendency of the network to have few nodes highly connected (Doncheva et al., 2012). This parameter is lower and higher in WT than both transgenic lines under dark and light conditions, respectively (Table S1), indicating that few nodes have high degree of connection in WT in the light. We next evaluated the appearance of new hubs and the preferential attachment in each genotype during dark-to-light transition. A hub-like node was considered as those with number of links above the average of the network under dark condition within each genotype. The number of hub-like nodes dropped from 22 to 3 in WT and from 25 to 17 in L3, whilst L13 increased from 28 to 37 after dark-to-light transition (Table S1). The preferential attachment was substantially different between the genotypes, in which WT, L3 and L13 have respectively 1, 9 and 26 nodes with higher number of links in the light than the average found in the respective genotype in the dark. Furthermore, the number of new hubs that appeared in the light is also higher in the transgenics, in which 2, 8 and 11 new hubs were found in WT, L3 and L13, respectively (Table S1).

### Integrating guard cell metabolomics data with stomatal speediness parameters

We next created a correlation-based network by combining guard cell metabolite profiling data with stomatal speediness parameters (*Sl*_max_ and t_50%_). We further integrated this with the previous multivariate analysis in order to obtain a systemic view of the physiological and metabolic alterations induced by *NtSUS2* that could potentially modulate stomatal speediness. Sixteen metabolites were found to be positively and negatively correlated to *Sl*_max_ and t_50%_, respectively (Figure 8A). Five of those metabolites (Ser, Ala, citrate, aconitate and mannitol) are also present in the VIP scores and S-plots of PLS-DA and orthoPLS-DA models carried out using relative guard cell metabolic changes during dark-to-light transition (Figure 8A). These results indicate that these metabolites are great contributors to the discrimination between the genotypes observed in both PLS-DA and orthoPLS-DA models and are positively correlated with the speed of light-induced stomatal opening. By contrast, trehalose and adipic acid were negatively and positively correlated to *Sl*_max_ and t_50%_, respectively (Figure 8A), indicating a negative correlation with the light-induced stomatal opening speediness. Interestingly, the metabolites that are positively and negatively related to the speed of light-induced stomatal opening have respectively lower and higher relative metabolic changes in both transgenic lines when compared to the WT during dark-to-light transition (Figure 8B). Comparing the dark-to-light transition in each genotype, it is interesting to note that aconitate increased significantly only in the WT, while adipic acid increase in all genotypes and mannitol decreased in L13 (Figure 8C). The higher accumulation of aconitate in WT and the lower relative changes in citrate observed in L3 further suggest that sucrose breakdown is important to feed the C6-branch of the TCA cycle. Collectively, our results suggest that *NtSUS2* is important for guard cell metabolism and, by consequence, for the regulation of light-induced stomatal opening and WPT.

**Figure 8.**
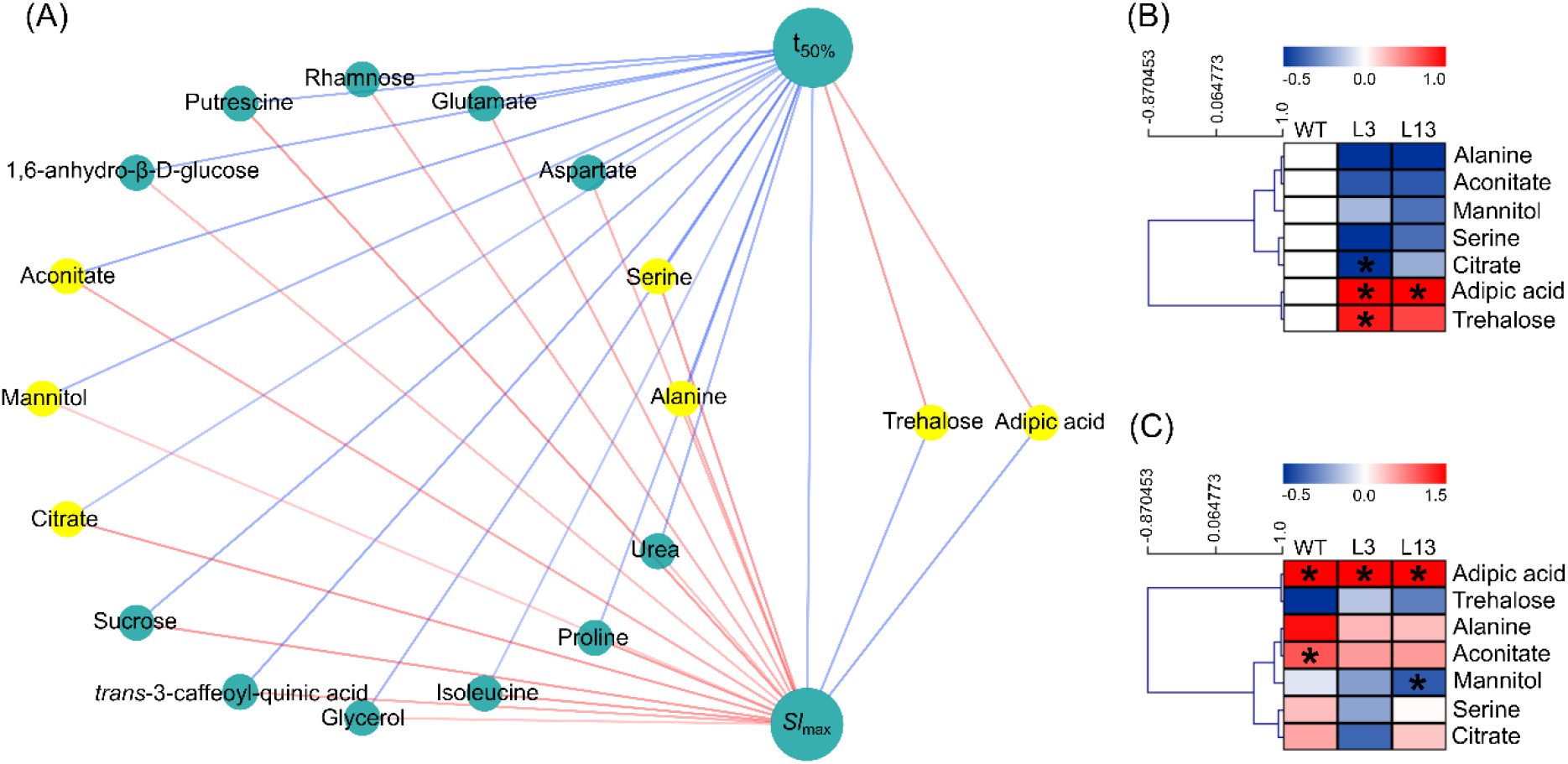
Integration of guard cell metabolic data with stomatal speediness parameters from *Nicotiana tabacum* L. wild type (WT) and transgenic lines (L3 and L13) antisense to *NtSUS2* grown under greenhouse and well-watered conditions. (A) Correlation-based network was created using relative guard cell metabolic changes after dark-to-light transition and maximum slope (*Sl*_max_) and half-time (t_50%_) needed to *g*_s_ reach steady state during light-induced stomatal opening. Yellow nodes highlight metabolites found in the S-plots of the orthoPLS-DA. Blue and red links indicate negative and positive correlation, respectively. (B) Heat map representation of the relative guard cell metabolic changes during dark-to-light transition, normalized according to the WT values and (C) relative guard cell metabolic contents in light, normalized by the dark values, of the yellow nodes of the figure A. Asterisks (*) indicate significant difference from WT by Student’s *t* test at 5% of probability (*P* < 0.05). (n = 5).

## Discussion

### Guard cell NtSUS2 is important for the regulation of whole plant transpiration

Sucrose has long been pointed out as an important metabolite that regulates stomatal movements (Granot & Kelly, 2019; Talbott & Zeiger 1998). Previous studies from our group highlight that manipulating the expression of *StSUS3* alter *g*_s_, with slight impacts on *A*, WPT and biomass production (Antunes et al., 2012; Daloso et al., 2016b). Here, we generated tobacco transgenic plants antisense for the *StSUS3* under control of the guard cell specific KST1 promoter. This led to reduced expression of ortholog *NtSUS2* in guard cell (Figure 1C) and decreased *g*_s_ in the transgenic lines (Figure 2E). These results are in agreement with those observed in potato plants expressing an antisense construct targeted against *SUS3* under 35S promoter (Antunes et al., 2012). However, it is important to highlight that the use of the constitutive 35S promoter reduced *A* up to ∼18%, whilst here the KST1-mediated reduction in guard cell *NtSUS2* expression have slightly reduced *g*_s_ with minor impact on *A* (∼10% in average) under well-watered conditions. This highlights the importance to use cell-specific promoters (Lawson et al., 2014), especially when the genetic manipulation involves sucrose metabolism, given its role in source-sink interaction.

Evidence linking sucrose metabolism and stomatal movement regulation comes from several genetic reverse studies. Transgenic plants with altered guard cell sucrolytic activity (Antunes et al., 2012; Daloso et al., 2016b; Ni, 2012), increased expression of guard cell hexokinase (Kelly et al., 2013, 2019; Lugassi et al., 2015), decreased expression of a guard cell plasma membrane sucrose (*SUT1*) and hexose (*STP*) transporters (Antunes et al., 2017; Flütsch, Nigro et al., 2020) or lacking key enzymes of starch breakdown (Flütsch et al., 2020b; Horrer et al., 2016) have altered *g*_s_ and/or stomatal aperture. These studies collectively indicate that sugar homeostasis and sucrose breakdown in guard cells are important to sustain the energetic and metabolites demand of guard cells during light-induced stomatal opening (Lawson & Matthews, 2020). They further support the idea that sucrose breakdown and hexose phosphorylation mediated by hexokinase is important during stomatal closure (Granot & Kelly, 2019). Although it seems contradictory, it is important to highlight that plants overexpressing hexokinase in guard cells can have higher or lower *E* depending on the environmental conditions (Lugassi et al., 2015). Similarly, the reductions in WPT observed in our transgenic lines were more prominent under fluctuating environmental (greenhouse) conditions, when compared to plants growing under controlled (growth chamber) conditions. Furthermore, WPT ranges between well-watered and water restriction periods, in which transgenic lines transpired less under well-watered and tend to transpire more under water shortage conditions. These results indicate that guard cell sucrose metabolism is important to regulate stomatal movements according to the prevailing environmental condition, highlighting the complexity and the plasticity of guard cell metabolism in responding to changes in the surrounding environment (Daloso et al., 2016a; Zeiger et al., 2002).

It is noteworthy that in the hottest and driest days of the experiment (days 4, 5 and 6 – Figure 3A), both L3 and L13 lost up to 44% less water than WT plants (Figures 3B,C). Furthermore, the L3 and L13 transgenic lines conserved 416 and 433 g H_2_O plant^-1^ through eleven days, respectively (Figure 3). Given that no changes in SD among the genotypes was observed (Figures S5A), this indicates that the lower WPT is not associated to changes in SD. Additionally, the WPT per leaf area remained significantly lower in L3 than WT at the days 5, 8, 9 and 10 of the experiment, in which no differences in leaf area between L3 and WT was observed at these days (Figures 3E-G). This indicates that smallest leaf area was also not the cause of the decreased WPT found in the transgenic lines. It seems likely therefore that the lower WPT of the transgenic lines is associated to the *NtSUS2*-mediated guard cell metabolic changes, as evidenced by both multivariate and metabolic network analyses (discussed below).

### On the role of guard cell metabolism for stomatal speediness regulation

The speediness of stomatal responses to environmental cues has recently received considerable attention to understand both the evolutionary origin of active stomatal control and its potential to maximize plant growth and WUE (Brodribb & McAdam, 2017; Flütsch et al., 2020a,b; Lima et al., 2019; Papanatsiou et al., 2019; Sussmilch, Roelfsema & Hedrich, 2019). However, the mechanisms regulating stomatal speediness remain far from clear (Lawson & Vialet-Chabrand, 2019). It is known that the degradation of starch, sucrose and lipids are important during light-induced stomatal opening (Daloso et al., 2015; Daloso et al., 2016b; Horrer et al., 2016; McLachlan et al., 2016). Given the role of malate as a counter ion of K^+^ during light-induced stomatal opening, it was long hypothesized that malate would be the fate of the carbon released from starch breakdown (Horrer et al. 2016; Lasceve, Leymarie & Vavasseur, 1997; Outlaw & Manchester, 1979; Schnabl, 1980; Schnabl, Elbert & Krämer, 1982; Talbott & Zeiger, 1993). However, recent evidence indicates that starch degradation and sugar homeostasis within guard cells are key to speed up light stomatal responses, but this was not associated to a differential malate accumulation among WT and *amy3 bam1* double mutant (Flütsch et al., 2020b). Additionally, no ^13^C enrichment in malate, fumarate and succinate was observed in guard cells under ^13^C-sucrose feeding during dark-to-light transition, but a substantial part of the ^13^C released from ^13^C-sucrose was incorporated into Gln under this condition (Medeiros et al., 2018). It seems likely therefore that the glycolytic fluxes are not directed to malate synthesis, as long hypothesized. In agreement with this idea, malate was not correlated to stomatal speediness and did not appear in the S-plots or VIP score lists here, despite the differences found in *g*_s_, stomatal speediness and WPT between WT and the transgenic lines. By contrast, other TCA cycle related metabolites such as citrate, aconitate, Asp, Glu and Pro were positively associated to stomatal speediness (Figure 8A) and citrate, aconitate, isocitrate and fumarate were included in the S-plots and the VIP score lists of at least one transgenic line (Figures 6 and S5).Furthermore, these metabolites have higher relative level in the WT when compared to both transgenic lines after dark-to-light transition (Figures 6 and S5), suggesting a higher activation of the TCA cycle and associated amino acid biosynthetic pathways in WT, which may contribute to explain the fast light stomatal response in this genotype.

Our results collectively further indicate that the decreased guard cell *NtSUS2* expression reduced the amount of substrate for glycolysis, which, in turn, affected the synthesis of amino acids. This idea is supported by the multivariate analyses in which Ala and Ser have lower accumulation in the transgenics and were included in both S-plots and VIP score lists (Figures 6 and S5). Furthermore, six amino acids (Ala, Ser, Asp, Glu, Ile and Pro) were positively correlated with stomatal speediness (Figure 8A). In contrast, trehalose and adipic acid were negatively associated to stomatal speediness (Figure 8A) and have higher relative level in the transgenics than the WT after dark-to-light transition (Figure 8B). Whilst information concerning the role of adipic acid in plant metabolism is scarce (Rodgman & Perfetti, 2013), trehalose metabolism has been closely associated to the regulation of stomatal movements (Daloso et al., 2016a; Figueroa & Lunn, 2016; Lunn et al., 2014). Although the exact mechanism by which trehalose metabolism contribute to regulate stomatal movements remain unclear, recent findings indicate that sugar homeostasis in guard cells is important to speed up light stomatal responses (Flütsch et al., 2020a,b) and to maximize the rapid early morning increase in *g*_s_ (Antunes et al., 2017). Thus, given that trehalose and trehalose-6-phosphate are closely associated to starch, sucrose, organic acids and amino acids metabolisms (dos Anjos et al., 2018; Figueroa et al., 2016; Martins et al., 2013), the differential accumulation of trehalose in the transgenics suggests that sugar homeostasis has been at least partially compromised in the transgenics. This idea is further supported by the fact that several sugars such as allose, idose, maltose, glucose, psicose and fructose were found in S-plots or VIP score lists in at least one transgenic line (Figures 6C,D; S5C,D). Sucrose was positively correlated with stomatal speediness (Figure 8A). Given that SUS works on both sucrose synthesis and degradation directions, this reinforce the idea that sugar homeostasis has been compromised in the transgenic lines, which in turn contributed to reduce the synthesis of organic acids and amino acids and the stomatal speediness in the transgenics in the light.

Our metabolic network analysis indicates that mild reductions in *NtSUS2* expression substantially alter both the topology and the connectivity of the guard cell metabolic network during dark-to-light transition. It seems likely therefore that *NtSUS2* is an important regulator of guard cell metabolism, highlighting why this gene is highly expressed in guard cells, when compared to mesophyll cells (Bates et al., 2012; Bauer et al., 2013; Yao et al., 2020). Given the complexity of guard cell metabolism, instead of using the classical reductionism perspective of searching for particular genes, proteins and metabolites that regulate stomatal speediness, a systemic view of the guard cell metabolic changes during dark-to-light transition might offer better possibilities to fully comprehend the regulation of stomatal kinetics (Medeiros et al., 2015). For this, incorporating dynamic metabolic data into genome scale metabolic models will certainly contribute to improve our understanding on the metabolism-mediated stomatal kinetics regulation. However, it is still unclear what are the key regulatory points in the guard cell metabolic fluxes. Evidence suggests that the fluxes toward glycolysis and the TCA cycle differ substantially in guard cells compared to mesophyll cells in the light (Hedrich, Raschke & Stitt, 1985; Horrer et al., 2016; Medeiros et al., 2018; Robaina-Estévez et al., 2017; Zhao & Assmann, 2011). This probably involves a different regulation of key glycolytic enzymes, which collectively may coordinate the fluxes from starch and sucrose breakdown toward the TCA cycle and associated pathways. However, diffuse information regarding the mechanisms that regulate guard cell metabolism hamper our understanding on the regulation of stomatal kinetics. Furthermore, guard cell metabolism complexity is often underestimated based on analogies made with mesophyll cells. However, guard cells have more than 1,000 genes that are differentially expressed with respect to mesophyll cells and seems to have higher plasticity in adjusting its metabolism according to soil water availability, light quality and intensity, CO_2_ concentration, air humidity and several others endogenous and environmental cues.

### Speeding up or slowing down stomatal opening? The route(s) toward WUE improvement

It has been estimated that slower stomatal opening can limit *A* by up to 10% (McAusland et al., 2016). Thus, increasing stomatal speediness through plant metabolic engineer could leads to increased crop yield and/or WUE (Lawson & Vialet-Chabrand, 2019). Indeed, optogenetic manipulation of a synthetic light-gated K^+^ channel (*BLINK1*) increased stomatal speediness to light and ultimately leads to increased biomass production without major impacts in water use by the plant (Papanatsiou et al., 2019). However, such engineered faster stomata could lead to unnecessary water losses in crops grown under extreme dry conditions. Thus, it is important to highlight that the strategies to improve crop WUE through genetic engineering should consider the water regime of the environment in which the plants will grow (Gago et al., 2014). In this context, our results highlight that slower stomatal responses and reduced *g*_s_ and WPT could also represent advantages for plants growing under fluctuating and dry conditions. An alternative possibility is thus to obtain stomata with faster closure and slower opening, in a scenario which *A* is not highly restricted by *g*_s_ during light imposition, *i*.*e*. the slower stomatal opening would not have substantial impacts on *A*, as observed in different angiosperms species (McAusland et al., 2016). Given that the mechanisms that regulate stomatal opening and closure differ substantially, it is possible to achieve this target through manipulation of the key regulators of both processes. However, several research has yet to be done to unveil the key regulators of stomatal speediness.

In conclusion, tobacco transgenic plants with antisense construction target to guard cell *NtSUS2* had slower stomatal opening in the light and decreased steady-state *g*_s_ and WPT under well-watered conditions. Furthermore, the transgenic lines showed higher or lower reduction in WPT under short water restriction periods, indicating a greater effective use of water under these conditions. Our results provide further evidence that *NtSUS2* is an important regulator of guard cell metabolic network and strengthen the idea that engineering guard cell metabolism is a promising strategy to decrease crop water consumption toward WUE improvement.

## Accession numbers

Sequence from *StSUS3* and *NtSUS1-7* genes data used on this article can be found in the <u>National Center for Biotechnology Information</u> databases (https://www.ncbi.nlm.nih.gov/) under accession numbers: AY205084.1, XM_016618980.1, XM_016595403.1, XM_016627888.1, XM_016610900.1, XM_016608012.1, XM_016585183.1, XM_016611295.1, respectively.

## Author contributions

F.B.S.F. and D.M.D. designed the research and experiments; F.B.S.F., R.L.G.B., R.S.C.B., S.A.C. and D.B.M. performed the experiments; all authors contributed to write the manuscript; D.M.D. obtained funding and is responsible for this article.

### Acknowledgments

This work was made possible through financial support from the National Council for Scientific and Technological Development (CNPq, Grant 428192/2018-1). We also thank the research fellowship granted by CNPq to D.M.D. and the scholarships granted by CNPq to F.B.S.F. and R.L.G.B. and the Brazilian Federal Agency for Support and Evaluation of Graduate Education (CAPES-Brazil) to R.S.C.B. and S.A.C.

