## Supplemental data for "Mild reductions in guard cell *sucrose synthase* 2 expression leads to slower stomatal opening and decreased whole plant transpiration in tobacco"

***Supplemental material***

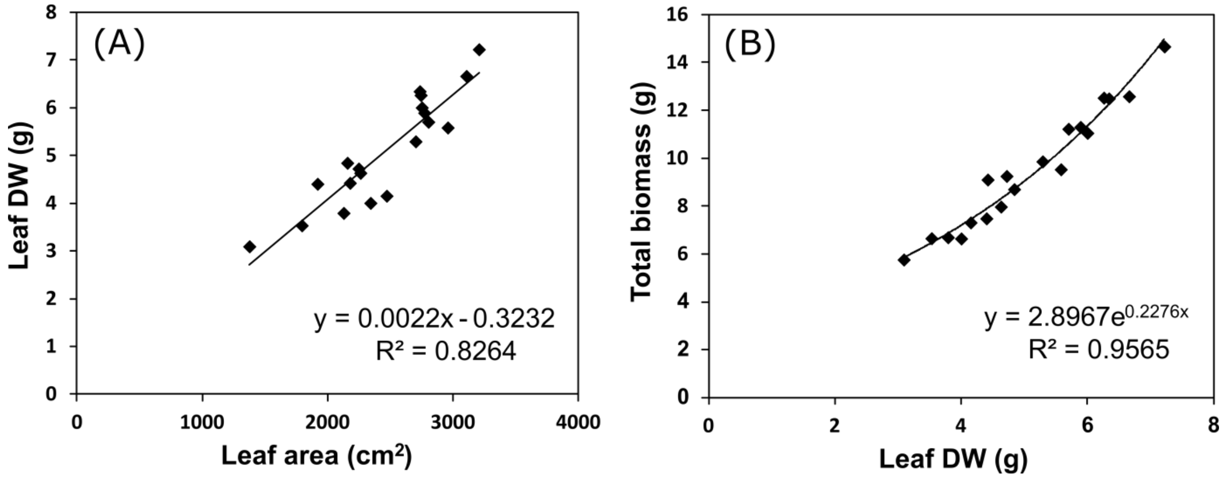


**Figure S1.** Regression analysis of growth parameters of *Nicotiana tabacum* L. wild type (WT) and transgenic lines (L3 and L13) antisense to *NtSUS2* grown under greenhouse and well-watered conditions. (A) Linear regression was carried out using leaf area and leaf dry weight (DW) to estimate leaf DW of the initial of the experiment. (B) After that, initial total biomass was estimated through the leaf DW values through exponential regression.

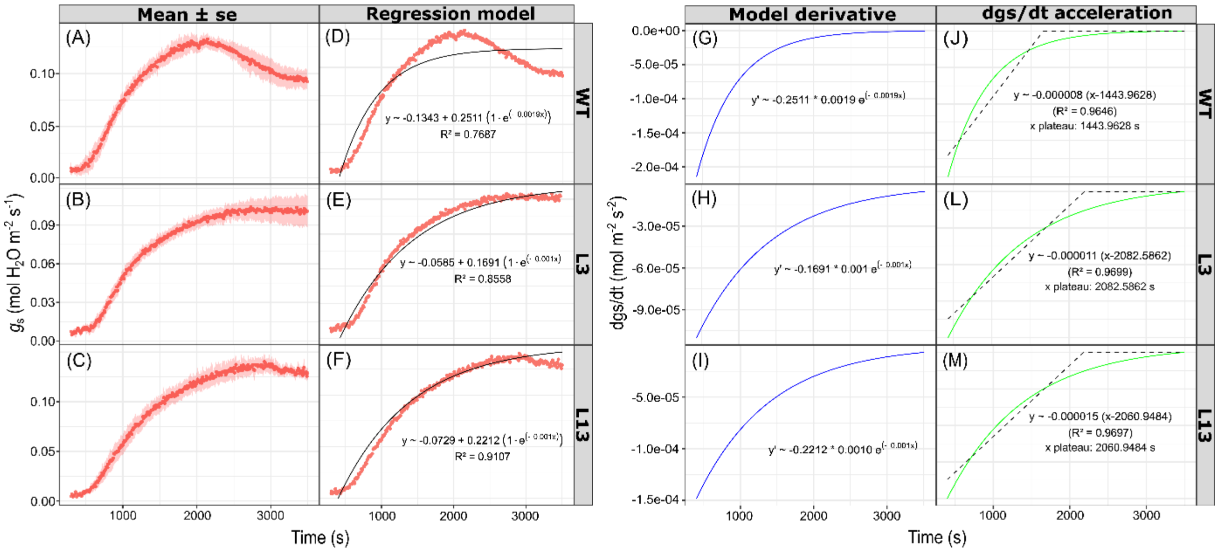


**Figure S2.** Stomatal conductance ($g_s$) responses (A-C) observed in *Nicotiana tabacum* L. wild type (WT) and transgenic lines (L3 and L13) antisense to *NtSUS2* grown under short water deficit stress during dark-to-light transition (0 to 1000 PPFD, in μmol $m^{-2}$ $s^{-1}$) (n = 3 ± SE). The total $g_s$ observed were best-fitted to non-linear model (D-E). By obtaining the equation model, the time derivatives were calculated (G-I) and further fit by linear plateau model (J-M) in order to estimate the time when $g_s$ reached maximum rate of change.

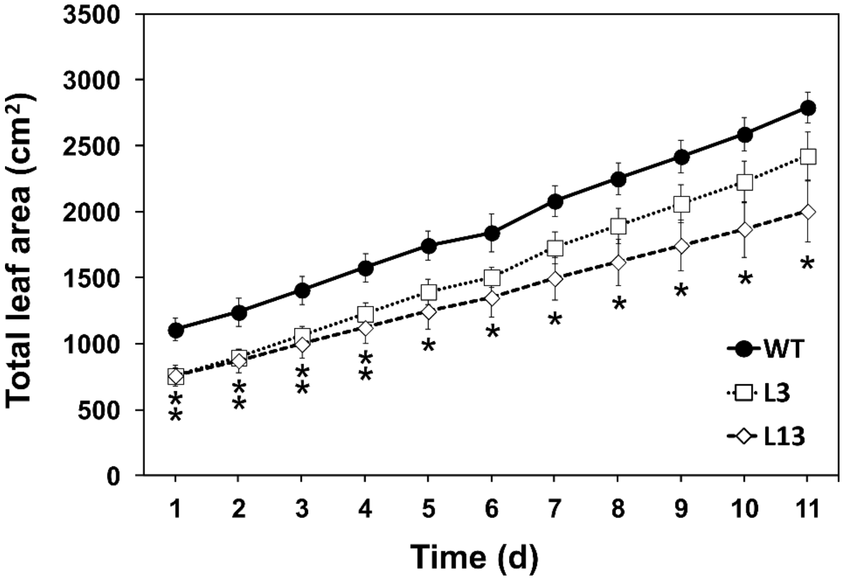


**Figure S3.** Total leaf area (LA) of *Nicotiana tabacum* L. wild type (WT) and transgenic lines (L3 and L13) antisense to *NtSUS2* grown under greenhouse and well-watered conditions. The measurements were taken during the whole plant transpiration experiment. LA was measured at the days 1, 5 and 11, whilst in the other days LA was estimated through linear regression analysis. One (*) and two asterisks (**) indicate that one or two transgenic lines are significant different from WT by Student's *t* test at 5% of probability ($P < 0.05$), respectively. (n = 5 ± SE).

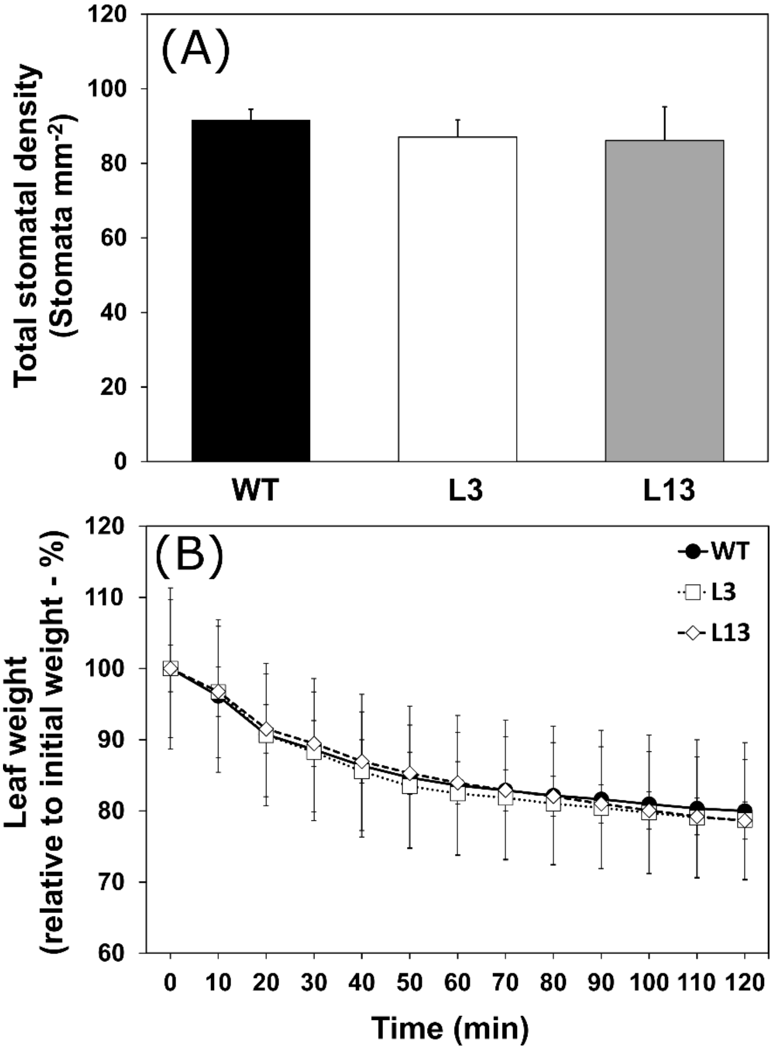


**Figure S4.** Stomatal density (A) and weight loss of detached leaves (B) of *Nicotiana tabacum* L. wild type (WT) and transgenic lines (L3 and L13) antisense to *NtSUS2* grown under greenhouse and well-watered conditions. Both measurements were taken in fully expanded leaves (n = 5 ± SE).

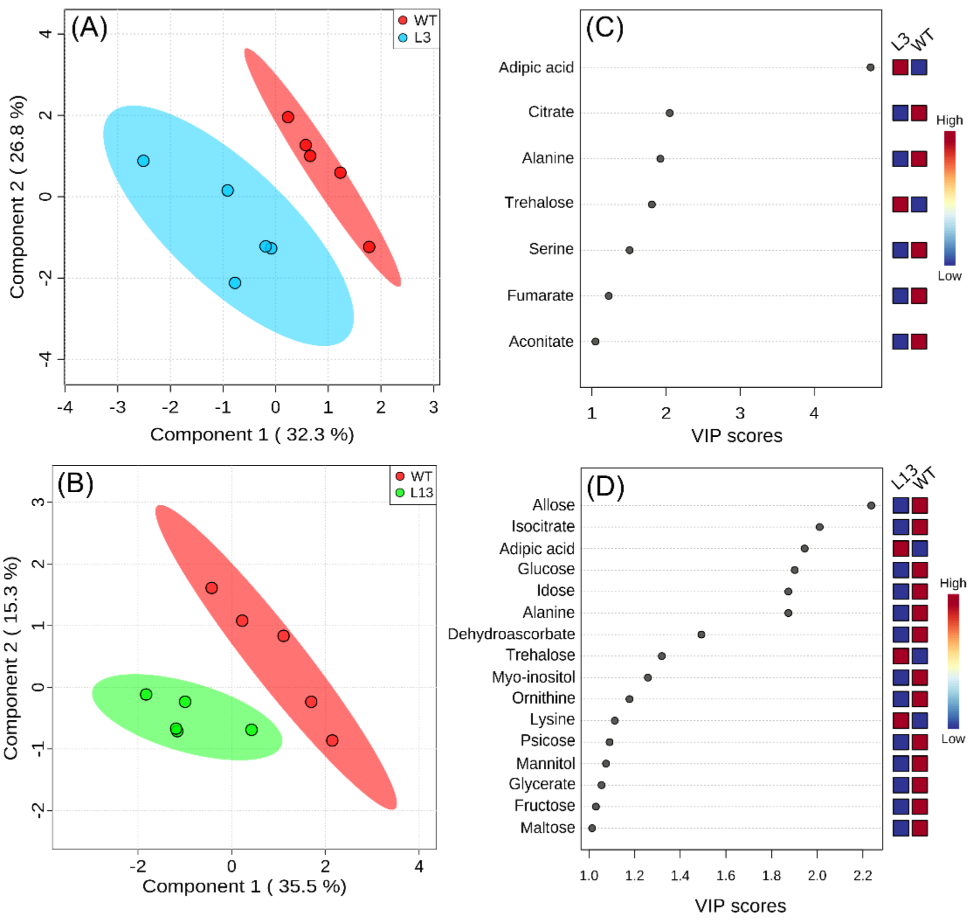


**Figure S5.** Partial least squares-discriminant analysis (PLS-DA) of guard cell metabolomics data. (A-B) WT *vs* transgenic lines PLS-DA. (C-D) Variable importance in projection (VIP) scores of the respective PLS-DA with metabolite list ranked from top to down for more important metabolites in PLS-DA models. These analyses were carried out comparing WT *vs* L3 (A, C) and WT *vs* L13 (B, D) separately using the relative guard cell metabolic changes observed in each genotype after dark-to-light transition. These analyses were performed using the MetaboAnalyst platform. (n = 5).

**Table S1.** Correlation-based metabolic network parameters from *Nicotiana tabacum* L. wild type (WT) and *NtSUS2* antisense transgenic (L3 and L13) plants grown under greenhouse and well-watered conditions. Data from relative guard cell metabolite content were used to create correlation-based metabolic networks for each genotype in the dark and after the transition to the light. These analyses were performed using CYTOSCAPE software. (n = 5).

| **Network parameter** | **Dark** | | | **Light** | | |
|---|---|---|---|---|---|---|
| | **WT** | **L3** | **L13** | **WT** | **L3** | **L13** |
| Clustering coefficient | 0.575 | 0.556* | 0.424* | 0.351 | 0.587** | 0.510** |
| Network centralization | 0.187 | 0.172* | 0.187 | 0.235 | 0.132* | 0.199* |
| Network density | 0.237 | 0.168* | 0.154* | 0.108 | 0.145** | 0.205** |
| Network heterogeneity | 0.338 | 0.464** | 0.668** | 0.838 | 0.447* | 0.523* |
| Connected components | 1.000 | 3.000** | 11.000** | 12.000 | 1.000* | 7.000* |
| Average number of links | 11.591 | 8.200* | 7.387* | 4.980 | 7.306** | 9.836** |
| Number of hub-like nodes | 22.000 | 25.000** | 28.000** | 3.000 | 17.000** | 37.000** |
| Preferential attachment | - | - | - | 1.000 | 9.000 | 26.000 |
| Hub appearance in the light | - | - | - | 2.000 | 8.000 | 11.000 |

*decreased relative to WT
**increased relative to WT
